# Identification of genes required for swarming motility in *Bacillus subtilis* using transposon mutagenesis and high-throughput sequencing (TnSeq)

**DOI:** 10.1101/2022.03.07.483392

**Authors:** Sandra Sanchez, Elizabeth V. Snider, Xindan Wang, Daniel B. Kearns

## Abstract

*Bacillus subtilis* exhibits swarming motility, a flagellar-mediated form of surface motility. Here we use high-throughput transposon mutagenesis and sequencing (TnSeq) to identify candidate genes required for swarming. The TnSeq approach identified all of the known genes required for flagellar biosynthesis as well as nearly all of the previously reported regulators that promote swarming. Moreover, we identified an additional 36 genes that improve swarming and validated them individually. Among these, two mutants with severe defects were recovered including *fliT*, required for flagellar biosynthesis, and a gene of unknown function *yolB*, whose defect could not be attributed to a lack of flagella. Our work indicates that TnSeq is a powerful approach for the identification of genes required for motility that drives outward expansion of a colony on plates.

**IMPORTANCE:** In TnSeq, transposons are randomly inserted throughout the chromosome at a population level, but insertions that disrupt genes of essential function cause strains that carry them to fall out of the population and appear under-represented at the sequence level. Here we apply TnSeq to the non-essential phenotype of motility in *B. subtilis* and spatially select for cells proficient in swarming. We find that insertions in nearly all genes previously identified as required for swarming are under-represented in TnSeq analysis, and we identify 36 additional genes that enhance swarming. We suggest that TnSeq is a powerful tool for the genetic analysis of motility and likely other non-lethal screens for which enrichment is strong.

## INTRODUCTION

The ancestral strain of *Bacillus subtilis* NCIB3610 is capable of swarming motility, an active and social form of flagellar-mediated movement that occurs atop a solid surface (1,2). Swarming over surfaces is thought to be different from swimming in liquid despite the fact that *B. subtilis* uses the same flagellar system for both behaviors. For example, when swimming cells are inoculated onto a soft agar surface, a lag period of immobility precedes the initiation of swarming suggesting that a physiological change is necessary for the behavior (1,3). One contributor to the lag is production of an extracellular surfactant that reduces surface tension and creates a thin layer of fluid within which to move (4,5). Another factor that contributes to the lag is the time it takes to increase flagellar biosynthesis and increase the flagellar density (3, 6,7). To determine the genetic requirements for swarming, a low-throughput transposon-based genetic screen was performed in which approximately 12,000 mutant colonies were picked, individually inoculated on miniature swarm agar plates, and manually screened for the inability to completely colonize the surface after overnight incubation (8). The forward genetic screen revealed a wide variety of “*fla*” mutants defective in flagellar biosynthesis as well as a mutant class called “*swr*” that abolished swarming motility but not swimming.

Study of swarming defective mutants has informed our understanding of how swarming and swimming differ. As one example, some mutants defective in the synthesis or activation of the extracellular lipopeptide surfactin abolished or impaired swarming, but these mutants could be rescued by providing purified surfactant exogenously (8-12). Other mutants exhibited cell autonomous defects related to the flagellum. As examples, cells mutated for the cytoplasmic proteins SwrA and EF-P decreased transcription and translation of flagellar genes respectively, while cells mutated for the membrane protein SwrB decreased the frequency of flagellar assembly (3,13-15). In addition, cells mutated for the putative efflux protein SwrC were impaired for flagellar function, apparently due to membrane accumulation of endogenously-synthesized surfactin (8,16). The original transposon screen was apparently not saturating, however, as other mutants have been discovered to have swarming defects like the two-component response regulator DegU and the polynucleotide phosphorylase PnpA that directly and indirectly activate flagellar gene expression, respectively (17-20). Thus, many of the genes that cell-autonomously promote swarming increase flagellar function or number.

*B. subtilis* synthesizes approximately 15-30 flagella per cell organized in a non-random pattern along the cell body (7,13). The biosynthesis of each flagellum involves highly conserved proteins that are assembled sequentially from the inside-out (21,22). First, a basal body containing a type III secretion system is assembled in the membrane with a gear-like rotor docked on the cytoplasmic surface. Next, the type III secretion system exports structural components for the axle-like rod and universal joint-like hook. Once the rod-hook is polymerized to a particular length, the secretion system changes specificity and exports components of the helical propeller-like filament. Finally, proton motive force-consuming stator channels associate with the flagellar rotor and create torque. *B. subtilis* encodes many but not all of the accepted flagellar structural proteins (22). Seemingly missing from the genome are homologs for the rod-cap protein that catalyzes rod polymerization and bushing proteins that surround the rod to stabilize flagellar rotation in the context of the envelope. Finally, while *B. subtilis* encodes many hydrolase enzymes that remodel peptidoglycan, there is currently no known hydrolase required to create space in the wall for rod assembly (23,24). Thus in *B. subtilis*, the “missing parts” are either unnecessary or await discovery.

Here we sought to identify additional genes required for flagellar assembly and/or swarming motility in *B. subtilis*. Previous work used transposon mutagenesis with a Tn10-based transposon somewhat prone to hotspot insertion, and manual screening of individual mutants was both cumbersome and limited by the diminishing return of the large set of genes already known to abolish motility when mutated (8). Here we revisit the forward genetic approach but use a mariner-based transposon for mutagenesis and screen with systems-level high-throughput transposon sequencing (TnSeq) (25-29). As proof-of-concept, almost all genes previously reported to be required for flagellar biosynthesis and swarming motility were under-represented in the TnSeq dataset. Moreover, individual testing of predicted candidates revealed thirty-six newly-identified mutants that exhibited swarming defects. All told, we show that TnSeq is a powerful approach to the genetic analysis of swarming motility. Moreover, the failure to reveal additional genes required for flagella structure in *B. subtilis* makes it increasingly likely that the “missing parts” either do not exist or are not required for flagella biosynthesis in this organism.

## RESULTS

### TnSeq analysis of swarming motility

Transposon mutagenesis and high-throughput sequencing (TnSeq) requires a large library of mutants with diverse insertion sites. We chose a mariner-based transposon for library generation because it has been found to insert randomly at TA sequences, a pattern that is enriched in the low G+C content bacterium *B. subtilis* (29-31). Based on preliminary mutagenesis using standard conditions and plating, approximately 7% of colony forming units were transposants. Using this information, we generated a control mutant library containing approximately 1 million colony forming units from which chromosomal DNA was purified, processed, and sequenced using an Illumina platform. Separately, this control library was inoculated to the center of swarm agar plates (**Fig. 1A**). When the swarm radius reached 25mm, the outer edge (greater than 15 mm off the center) was harvested by swabbing, and cells from 10 plates were combined to form a motility-enriched pool (**Fig 1A**). Five different motility-enriched pools were generated from which chromosomal DNA was purified and sequenced.

**Figure 1:**
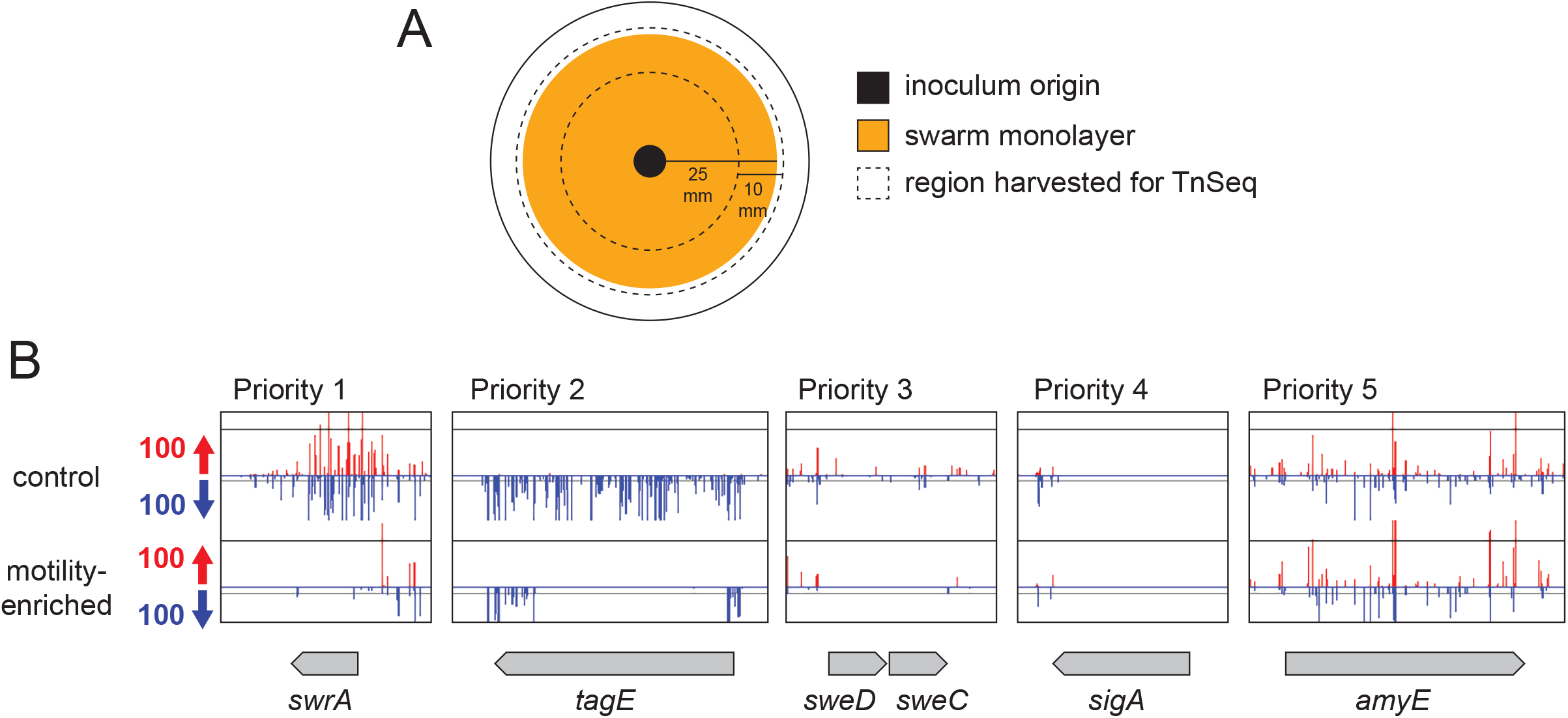
TnSeq priority classification of candidate genes defective in swarming motility. A) Top view diagram of a Petri plate used in the TnSeq selection for swarming motility. Outer circle represents perimeter of a petri plate. Black circle indicates the origin where the transposon mutagenized pool was inoculated. Orange circle indicates the area of the swarm with ∼25 mm radius. The region between the dotted lines indicate the region from each plate that was harvested for pooling the cells prior to DNA isolation and transposon sequencing. B) Sample genes with transposon insertion density graph generated using Artemis software. Transposon insertions oriented such that the spectinomycin resistance cassette is co-oriented with the inserted gene colored red, while insertions that counter-orient the resistance gene are colored blue. Length of the bars indicates the relative number of redundant insertions at that position. Gene length and orientation indicated as a gray arrow. Top dataset is from the control library, and bottom data set is from one of the motility-enriched pools. Priority 1 represented genes that passed mathematical criteria for differential insertion and the difference was readily visible on manual inspection. Priority 2 were those genes that failed at least one of the mathematical criteria but were rescued by a clear insertion differential by manual inspection. Priority 3 genes were those that passed mathematical criteria but did not appear to have a dramatic insertion differential by manual inspection. Priority 4 genes were those genes that passed mathematical criteria but were discarded for having few, if any, transposon insertions. Priority 5 genes were those genes that passed neither mathematical nor manual inspection criteria and were discarded.

An average of 14 million reads were obtained for both the control and motility-enriched samples. The complexity of a library was determined by comparing the number of possible TA sequence *mariner*-insertion sites in the chromosome relative to the number of TA sites that actually experienced transposon insertion in the pool. The *B. subtilis* wild type genome has 218,028 possible TA sites and the control library had at least two insertions in 53% of the TA sites, and at least 10 insertions in 40% of the TA sites. Thus, the control library sampled insertions that were both diverse and exhibited high redundancy. Next, the insertions sequenced from the motility-enriched pools were compared to the control dataset. We hypothesized that candidate genes required for swarming motility would be those genes that experienced a reduction in transposon insertion density specifically after selection for swarming proficiency.

To identify candidate genes worthy of further consideration, the transposon insertion density per open reading frame was compared by a combination of both mathematical and subjective criteria. Genes were mathematically included as potential candidates if they contained at least 10 potential TA sites within the coding region (i.e. weren’t so small that TA sites were inherently rare) and they experienced a statistically significant (P value <0.05) 10-fold reduction in representation after swarming selection for at least 3 out of the 5 swarming selected transposon mutant pools. Genes were also subjectively curated by manually scanning of the graphical representation of transposon peak insertion density using Artemis software (32). Manual scanning could either increase confidence in the mathematical inclusion of a particular gene or add additional genes as candidates if those genes did not meet the strict mathematical criteria but nonetheless appeared to have a visually conspicuous reduction in insertion density after motility-enrichment. Lastly, manual curation also excluded some mathematically-supported candidates if the number of insertions in that particular gene appeared to be low such that loss of those insertions during swarming could either be stochastic or due to a compounding growth defect. After the combined mathematical and manual screening of the data, genes were assigned a priority score from 1 to 5, with 1 being the highest priority and 5 being the lowest (**Fig 1B**).

Genes were given a priority score of 1 if they passed all mathematical criteria and were also supported by manual screening with an insertion differential that was noticeable by eye. Genes were given a priority score 2 if they failed one or more of the mathematical criteria but were rescued by manual screening because there was a visually dramatic reduction in insertions sites after motility-enrichment. Genes were given a priority score of 3 if they passed the mathematical criteria but manual screening indicated that the insertion density differential did not to be appear dramatic, often because the insertion density in the control condition was low. Genes were given a priority score of 4 if they passed the mathematical criteria but were excluded by manual analysis for having too few insertions under both control and motility-selected conditions. Finally, genes were given a priority score of 5 if they failed to meet both the mathematical criteria and failed to be rescued by manual screening. Only those genes with priority scores of 1-3 were considered worthy of further analysis and individual phenotypic testing for swarming motility.

To determine whether the TnSeq screen for swarming motility-defective mutants was functioning as expected, we analyzed the list of candidates for genes known to be required for flagellar assembly and function (**Table 1**). Many of the genes required for the early stages of flagellar assembly including the flagellar basal body, rod and hook, are encoded in the 32 gene *fla/che* operon and the insertion density in this region was reduced in the motility-enriched population (**Fig 2A**) (33-35). The genes dedicated to late-stage flagellar assembly, including the flagellar filament, are encoded in multiple cistrons in a region called the “second flagellar cluster” encoded 180° from the *fla/che* operon on the circular chromosome, and structural genes encoded in this region also experienced a reduction in insertion density (**Fig 2B**). Two other loci are required for flagellar assembly and/or function: the *flhO-flhP* operon encoding the distal rod proteins FlhO and FlhP, and the *motA-motB* operon encoding the force generating MotA-MotB stator units, and these operons also experienced a reduced insertion density (36-39). We conclude that the TnSeq screen for swarming-essential genes recovered every gene previously identified as being required for flagellar structure and function.

**Table 1:** Genes predicted by TnSeq that were previously shown to be required for swarming motility.

| Gene | PS<sup>1</sup> | Category<sup>2</sup> | Annotation<sup>3</sup> | Ref |
| --- | --- | --- | --- | --- |
| *cheA* | 1 | motility | chemotaxis, run generator | 1 |
| *cheB* | 2 | motility | *fla/che* operon, polar | 8 |
| *cheC* | 1 | motility | *fla/che* operon, polar | 8 |
| *cheD* | 1 | motility | *fla/che* operon, polar | 8 |
| *cheW* | 1 | motility | *fla/che* operon, polar | 8 |
| *cheY* | 1 | motility | chemotaxis, run generator | 1 |
| *clpC* | 3 | regulation | AAA+ protease, unknown | 7 |
| *clpX* | 3 | regulation | AAA+ protease, unknown | 7 |
| *cwlO* | 1 | envelope | PG hydrolase | 24 |
| *cwlQ* | 2 | envelope | PG hydrolase | 24 |
| *degU* | 1 | regulation | flagellar regulation | 18 |
| *efp* | 3 | translation | elongation factor P | 8 |
| *flgB* | 1 | motility | flagellar structure | 60 |
| *flgC* | 1 | motility | flagellar structure | 60 |
| *flgD* | 1 | motility | flagellar structure | 36 |
| *flgE* | 1 | motility | flagellar structure | 36 |
| *flgK* | 1 | motility | flagellar structure | 38 |
| *flgL* | 1 | motility | flagellar structure | 38 |
| *flgM* | 1 | motility | flagellar regulation | 60 |
| *flgN* | 1 | motility | flagellar structure | 61 |
| *flhA* | 1 | motility | flagellar assembly | 60 |
| *flhB* | 1 | motility | flagellar assembly | 60 |
| *flhF* | 1 | motility | flagellar regulation | 13 |
| *flhG* | 1 | motility | flagellar regulation | 13 |
| *flhO* | 1 | motility | flagellar structure | 36 |
| *flhP* | 1 | motility | flagellar structure | 36 |
| *fliD* | 1 | motility | flagellar structure | 62 |
| *fliE* | 1 | motility | flagellar structure | 60 |
| *fliF* | 1 | motility | flagellar structure | 60 |
| *fliG* | 1 | motility | flagellar structure | 60 |
| *fliH* | 1 | motility | flagellar assembly | 60 |
| *fliI* | 1 | motility | flagellar assembly | 60 |
| *fliJ* | 1 | motility | flagellar assembly | 60 |
| *fliK* | 1 | motility | flagellar regulation | 36 |
| *fliL* | 1 | motility | flagellar function | 60 |
| *fliM* | 1 | motility | flagellar structure | 60 |
| *fliO* | 1 | motility | flagellar assembly | 60 |
| *fliP* | 1 | motility | flagellar assembly | 60 |
| *fliQ* | 1 | motility | flagellar assembly | 60 |
| *fliR* | 1 | motility | flagellar assembly | 60 |
| *fliS* | 1 | motility | flagellar assembly | 62 |
| *fliY* | 1 | motility | flagellar structure | 60 |
| *gsaB* | 1 | translation | EF-P modification | 45 |
| *hag* | 1 | motility | flagellar structure | 1 |
| *minJ* | 2 | division | FtsZ localization factor | 63 |
| *motA* | 1 | motility | flagellar function | 37 |
| *motB* | 1 | motility | flagellar function | 37 |
| *pdeH* | 1 | regulation | flagellar function | 64 |
| *pnpA* | 1 | regulation | flagellar regulation | 19 |
| *rho* | 3 | transcription | transcriptional terminator | 65 |
| *sigD* | 1 | motility | flagellar regulation | 66 |
| *swrA* | 1 | motility | flagellar regulation | 8 |
| *swrB* | 1 | motility | flagellar assembly | 8 |
| *swrC* | 1 | motility | surfactin resistance | 8 |
| *swrD* | 1 | motility | flagellar function | 56 |
| *yabR* | 2 | unknown | unknown | 8 |
| *ylxF* | 1 | motility | *fla/che* operon, polarity | 60 |
| *ymfl* | 1 | translation | EF-P activation | 44 |
| *ynbB* | 1 | translation | EF-P modification | 45 |

<sup>1</sup>Each gene was assigned a priority score (PS) based on data analysis. A score of “3” indicates that the gene passed the criteria of a ratio of 0.1 or lower and a statistical P value of <0.05 and could not be eliminated by manual evaluation of the data. A score of “2” indicates that the gene did not pass the ratio cutoff but was rescued because it appeared to have an insertion differential by visual scanning of the data set. A score of “1” indicates that the gene passed the quantitative criteria and also showed strong insertion differential by visual scanning of the data set.
<sup>2</sup>Category is a general category of function assigned to the presumed or known function.
<sup>3</sup>Annotation is a brief specific description of the known or predicted function of the gene product.

**Figure 2:**
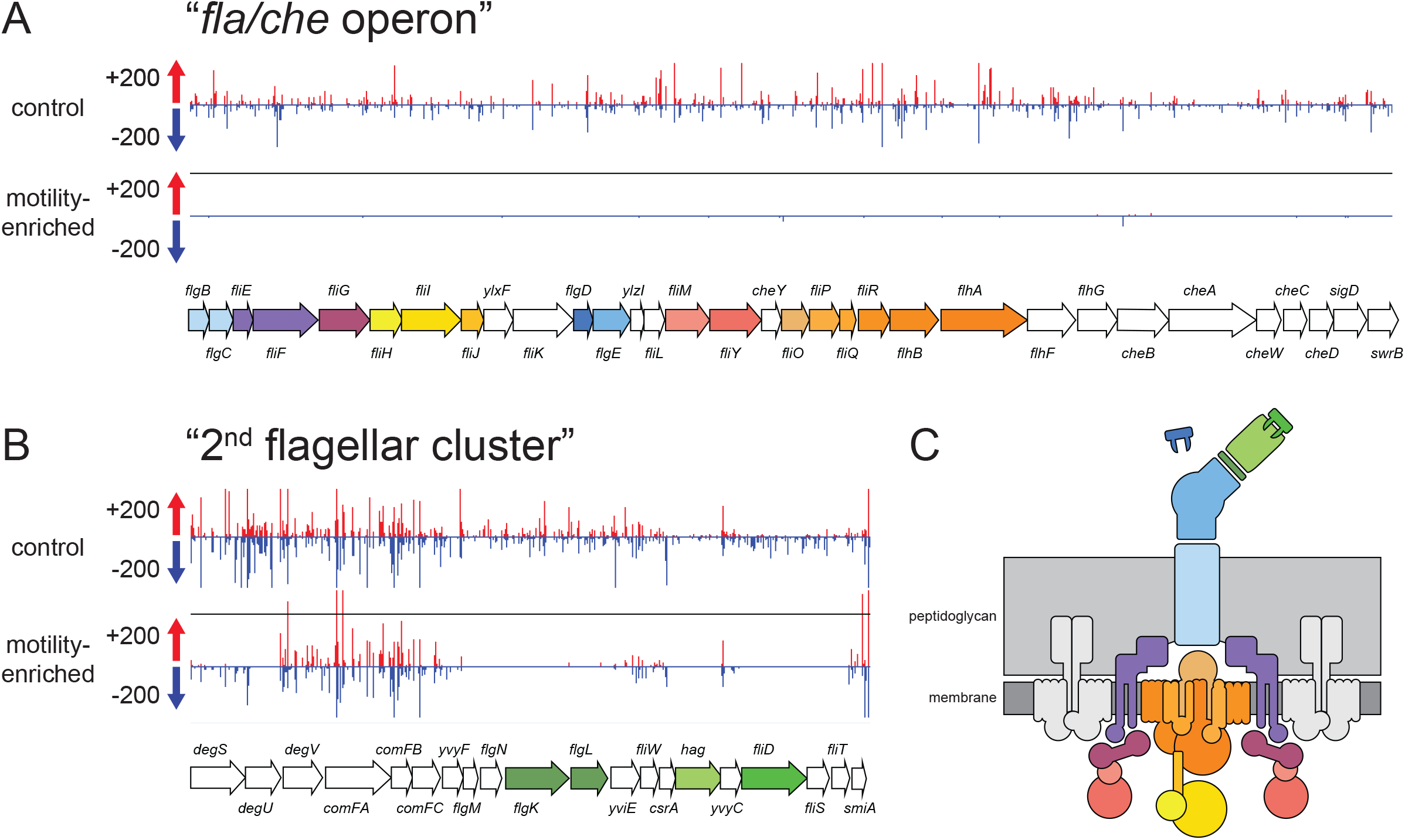
TnSeq identified flagellar structural genes necessary for swarming motility. Transposon insertion densities within A) the *fla/che* operon and B) the second flagellar cluster graphed by Artemis software. Transposon insertions oriented such that the spectinomycin resistance cassette is co-oriented with the inserted gene colored red, while insertions that counter-orient the resistance gene blue colored red. Length of the bars indicates the relative number of redundant insertions at that position. Gene length and orientation indicated as an arrow. Colored arrows indicate flagellar structural genes and the colors match the relative location in the flagellum as indicated in panel C. C) Cartoon diagram of the Gram-positive flagellum with parts color-coded to the match the genes that make the corresponding structure in panels A and B.

Our TnSeq screen also identified a number of regulatory genes that have previously been reported to specifically confer defects in swarming motility (**Table 1**). For example, reduced insertion density was observed in the genes encoding proteins necessary for increased flagellar number during swarming including the response regulator DegU, the master flagellar activator SwrA, and the activator of type III secretion SwrB (14,40-42). Genes encoding other proteins known to be strictly required for swarming such as the protease subunit ClpX, polynucleotide phosphorylase PnpA, and the putative surfactin efflux protein SwrC also experienced reduced insertion (7,8,19,20). Finally, the gene encoding Efp that enhances translation of a limiting flagellar structural protein, and genes encoding YmfI and GsaB involved in Efp activation were also under-represented or absent (15,43-45). We conclude that the TnSeq screen was successful at predicting regulators previously demonstrated to be required for swarming.

Some genes required for swarming motility were not predicted by the TnSeq analysis. For instance, the long *srfAA-srfAD* operon required for synthesis of the swarming surfactant, surfactin, the Com quorum sensing system that activates *srfAA-srfAD* gene expression, and the *sfp* gene required for enzymatic activation of the surfactin synthase complex all did not appear to be required (10,46,47). Their absence from the candidate list is likely explained because each gene contributes to the production of surfactin, and surfactin defects can be extracellularly complemented for swarming motility (1,48). Presumably, the other members of the mixed mutant pool provided surfactin sufficient for the rare deficient mutants. One other gene known to be required for swarming, which encodes the biofilm matrix and flagellar clutch repressor SinR, was not predicted to be required by the TnSeq analysis but this was likely because *sinR* experienced a low overall insertion density in the control library (49,50). The failure to obtain insertions in the *sinR* gene may have been stochastic or because *sinR* mutants de-repress biofilm matrix genes and excessive cell clumping may have impaired DNA recovery. All told, the TnSeq screen provided proof-of-principle results by predicting 59 out of 60 genes previously reported to be required for swarming in a cell-autonomous manner.

### TnSeq identifies new genes required for swarming motility

In addition to identifying the known motility genes, TnSeq analysis predicted 78 new candidate genes (**Table S1**), with priority scores from 1-3 (**Fig. 1B**). To validate these candidates, we made disruptions of those genes and quantitatively assayed the mutant strains for swarming motility. While many had behavior indistinguishable from wild type (**Table S1**), thirty-six of the mutants exhibited a defect in swarming motility (**Table 2**). Twelve of the mutants were phenotypically classified as “slow” because the defect in swarming motility appeared to be due to a reduced rate of colony expansion (**Fig 3, cyan**). Fifteen of the mutants were phenotypically classified as “extended lag” because the rate of swarm expansion appeared to be comparable to that of wild type but the period of immotility prior to swarming initiation was prolonged (**Fig 3, purple**). Seven of the mutants were phenotypically classified as “terracing” because they exhibited a periodic cessation and re-initiation of swarming motility that gave rise to a terraced appearance of the colony after overnight incubation (**Fig 3, orange**). Finally, two of the mutants were phenotypically classified as “severe” because they failed to expand substantially from the inoculation origin during the period of measurement and even after overnight incubation (**Fig 3, red**). We conclude that the TnSeq approach was successful at identifying new genes involved in swarming motility.

**Table 2:**
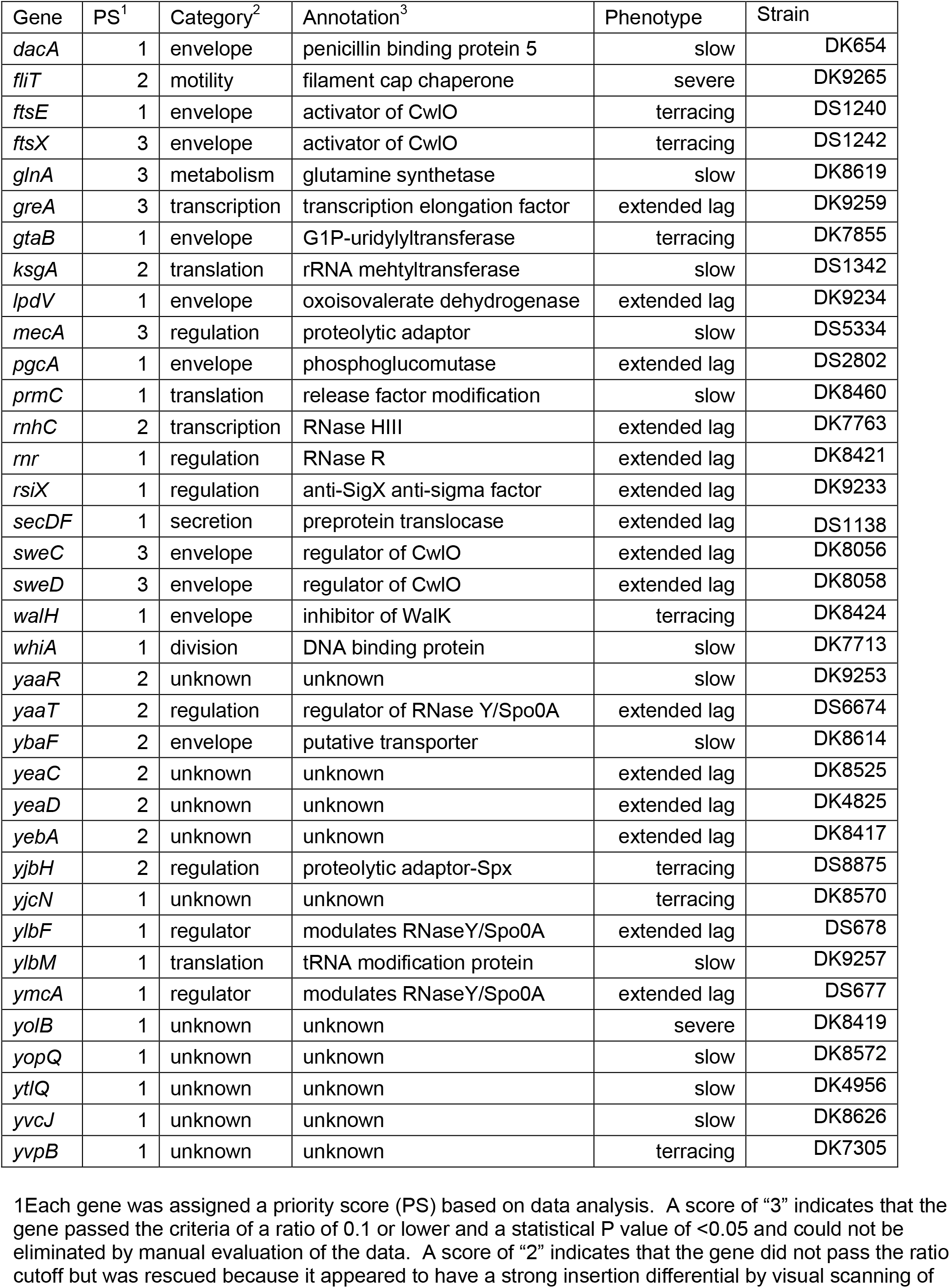
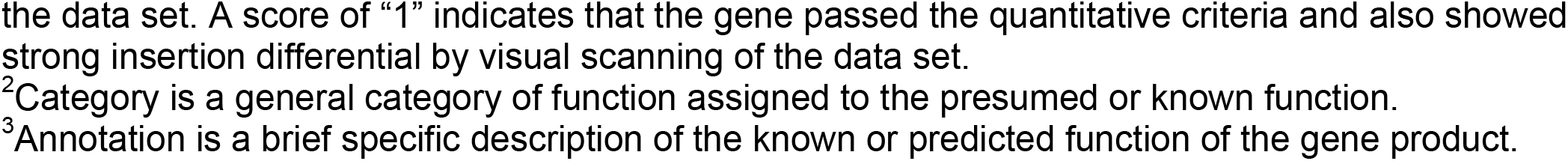
Candidate genes which when mutated conferred a defect in swarming motility.

**Figure 3:**
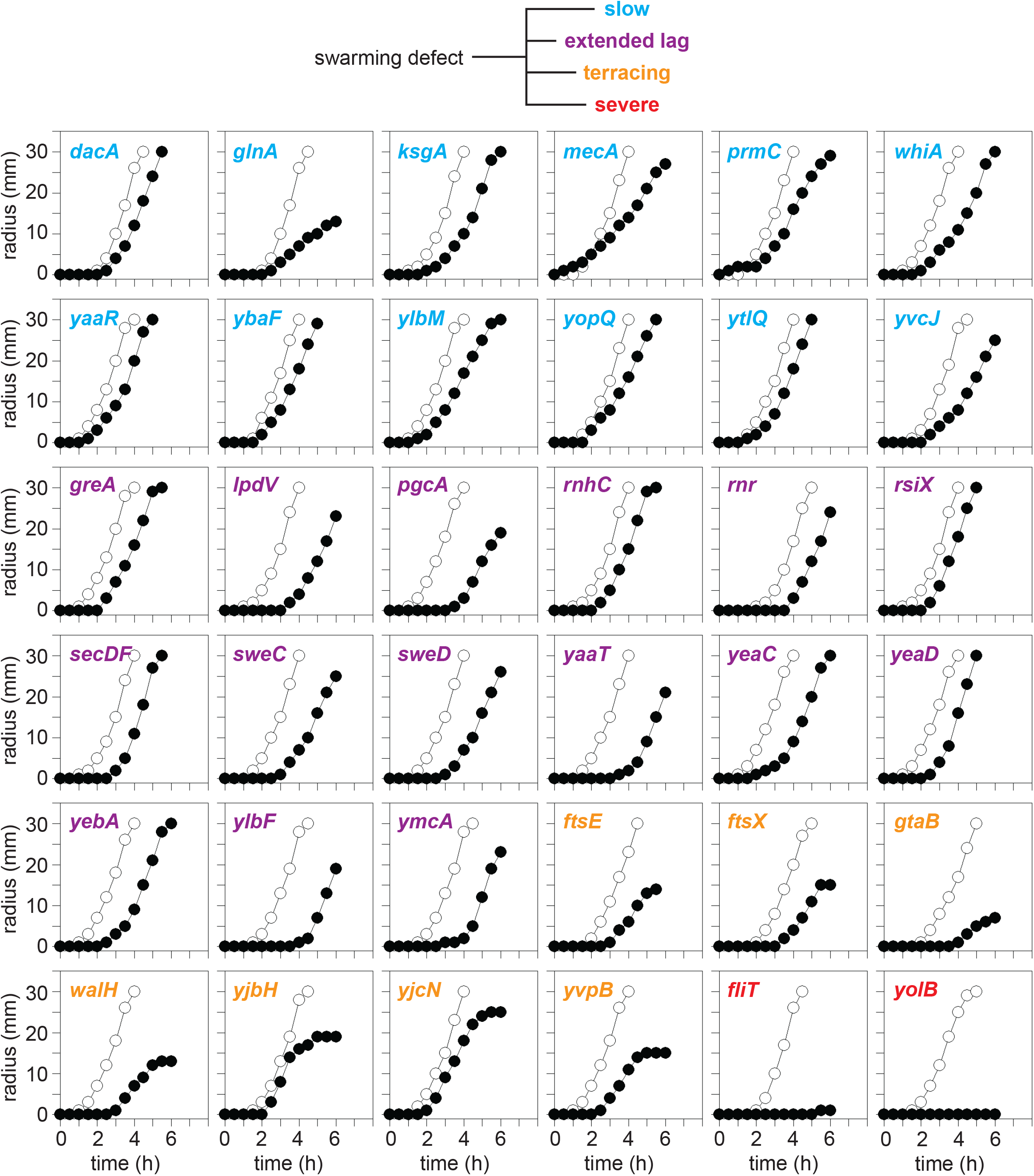
Newly-identified genes which when mutated exhibit defects in swarming motility. Quantitative swarm expansion assays of wild type (open circles) and mutant (closed circles) as indicated by the gene name in the upper right hand corner of the graph. Some wild type data has been duplicated as the same wild type data was used as a control for mutants that were analyzed for swarming concurrently. Each data point is the average of three replicates. Swarm graphs grouped according to their respective class of defect: cyan, slow swarm expansion; purple, extended lag period, orange, terracing mutants; red, severe swarm defect.

We noted that some of the mutants grew slowly in liquid media and we wondered whether growth defects, if any, were correlated with impaired swarming. Each mutant with a swarming defect was grown in a microtiter plate reader in triplicate, and the maximum growth rate during exponential growth was calculated. Relatively few mutants exhibited a dramatic decrease in growth rate relative to the wild type (**Fig 4**). Reduction in growth rate however, was more common in those mutants that exhibited a slow swarming phenotype. We infer that a reduction in growth rate is potentially one way to cause a defect in swarming motility but whether the swarming defect is directly due to the growth rate change or some other consequence of the mutation is unknown. Interestingly, mutants with reduced growth rate did not appear to increase the lag time prior to swarming (e.g. *glnA*) suggesting that growth rate and lag period were unrelated (**Fig 4**). Finally, we note that the mutants that conferred a severe defect in swarming motility, *fliT* and *yolB*, exhibited growth rates comparable to the wild type and we infer the swarming defect was due to some other factor.

**Figure 4:**
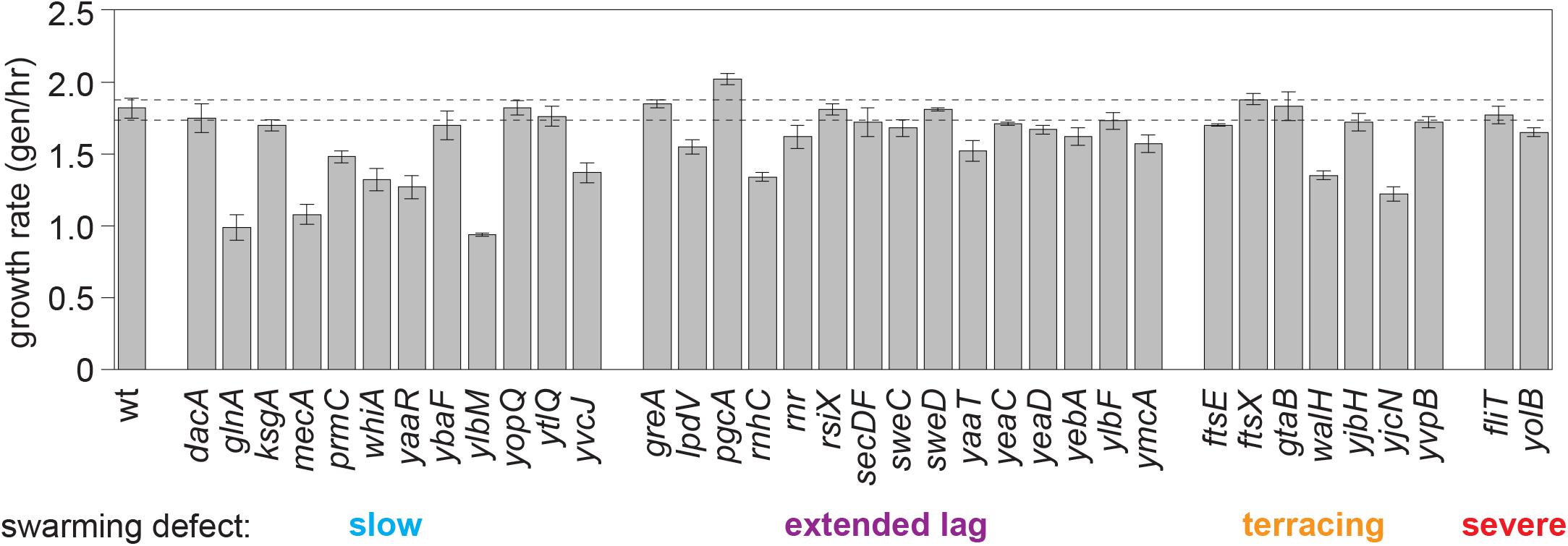
Growth rates of mutants with swarming defects. Cells were grown in a 96 well microtiter plate and rates are the average of three replicate experiments. Error bars are standard deviation. Dashed lines are extensions of the standard deviation of wild type for quick comparison.

One way in which a mutation can result in a severe defect in swarming motility is if the mutant is defective in flagellar biosynthesis. To detect flagellar biosynthesis, a version of the flagellar filament protein flagellin that can be fluorescently labeled with a maleimide derivative (*hag*^*T209C*^) (50) was introduced into both the *fliT* and *yolB* mutant backgrounds. Whereas wild type was proficient for flagella biosynthesis, the *fliT* mutant was defective (**Fig 5**). The *fliT* gene encodes a homolog of FliT, a bifunctional protein in *Salmonella* that primarily serves as the secretion chaperone for the flagellar cap protein FliD (51-55). The *fliT* flagellar filament defect observed in *B. subtilis* is consistent with a FliD secretion defect, and FliT is likely the last flagellar protein homolog that could have been predicted to have a motility defect by a reverse genetic approach. In contrast to *fliT*, the *yolB* mutant was proficient in flagellar biosynthesis (**Fig 5**). The *yolB* gene encodes the protein of unknown function YolB, and the reason a *yolB* mutant is defective in swarming is unknown. As flagella are required for swarming motility, we infer that all of the other swarming mutants with partial defects in behavior are flagella proficient. Thus, the TnSeq screen identified new genes involved in swarming motility but seemingly no additional *B. subtilis*-specific genes required for flagellar assembly.

**Figure 5:**
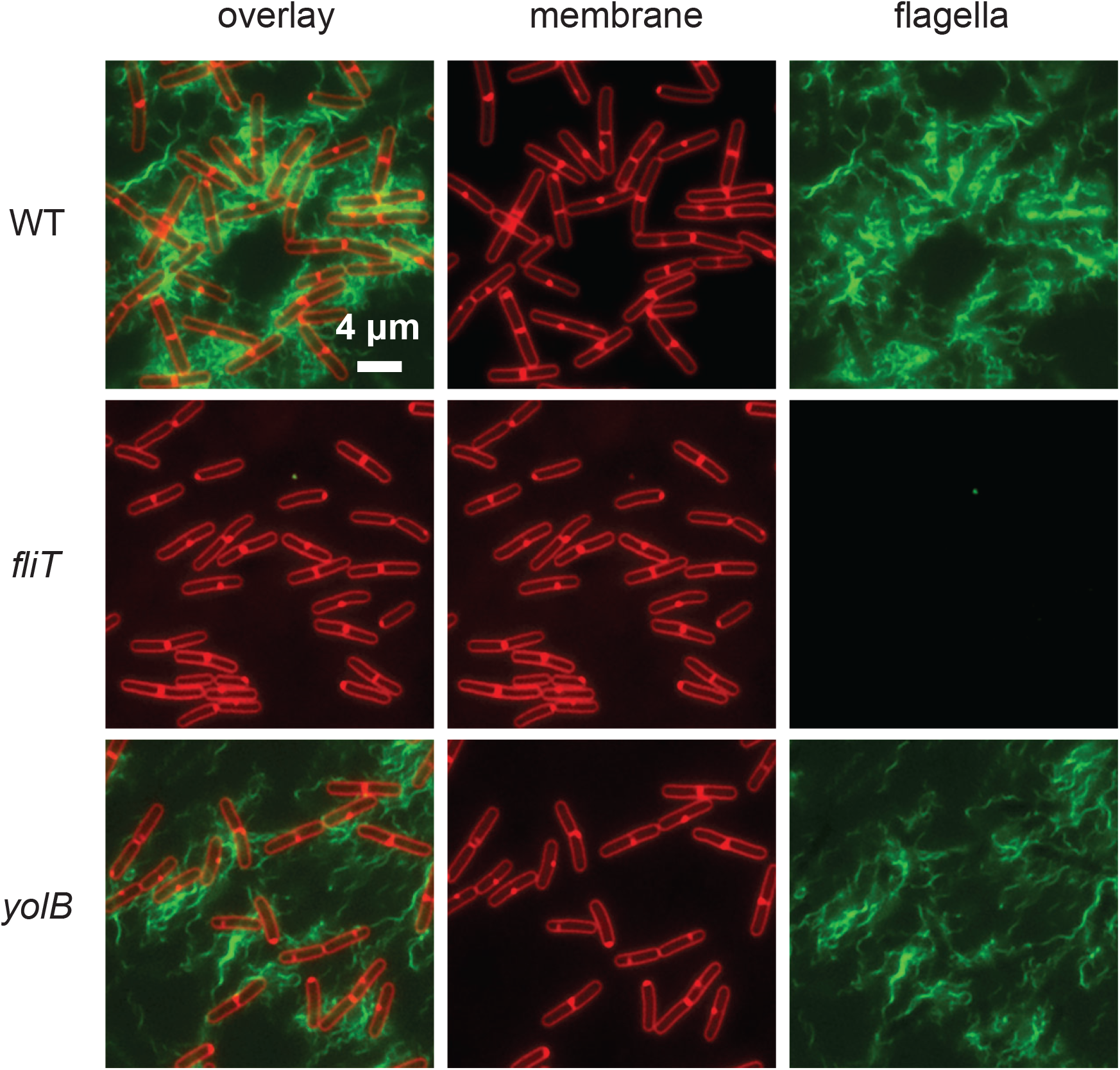
Cells mutated for *yolB* are proficient for flagellar biosynthesis. Fluorescent micrographs of WT (DS1916), *fliT* (DK9348), and *yolB* (DK8486) mutants expressing the flagellar filament allele Hag^T209C^ and stained for membrane with FM 4-64 (false colored red) and flagella with alexa fluor maleimide (false colored green).

## DISCUSSION

The high-throughput systems-level TnSeq protocol described here was highly successful in identifying genes required for swarming motility. By selecting the subpopulation of the transposon mutant pool that was swarming-proficient, individuals with transposon insertions in swarming-essential genes were excluded, and such genes experienced a reduction in insertion density after sequencing. Ultimately, our mathematical and manual screening of the insertion data was able to predict 59 out of 60 genes previously reported to be required for swarming. Moreover, the data indicated another 78 genes experienced a similar reduction in insertion density after motility enrichment and these became candidate genes for individual validation. Manual testing of individual mutants found that roughly half of the candidates exhibited a swarming defect when tested in isolation, proving that the approach could identify new genes involved in the phenotype. Moreover, we found that many of the mutants that reduced swarming were associated with either envelope maintenance or efficient translation, perhaps consistent with the need for high numbers of proteinaceous trans-envelope flagella. Finally, mutation of one gene *yolB*, resulted in a severe defect in swarming, consistent with being a new member of the “*swr*” genetic class.

The “*swr*” genetic class is defined as those genes which when mutated, abolish swarming motility but not swimming (Kearns et al., 2004). Moreover, the swarming defect of a *swr* mutant is cell autonomous and cannot be complemented extracellularly, thus excluding genes producing shared secreted products like surfactants and quorum signals. Preliminarily, *yolB* meets the criteria of a *swr* gene as mutation results in a severe swarming defect both in mixed mutant pools and in isolation, while seemingly not impairing the production of flagella or impairing swimming by wet mount observation. The *yolB* gene encodes YolB, a 118 amino acid protein of unknown function that has no domains conserved with other proteins in the database. How YolB might promote swarming motility is unclear. Perhaps it is needed to increase flagellar number on surfaces like SwrA, or increases the power to individual flagellar motors like SwrD (13,56). Whatever the case, YolB is encoded within the *B. subtilis* chromosomal prophage Spβ, and its activity may represent a form phenotypic conversion by a horizontally transferred element.

Another class of mutants that causes a severe defect in swarming motility is the *fla* class, whose mutation abolishes flagellar biosynthesis (1). Only one new gene, *fliT*, met the criteria for a *fla* class mutant. While *fliT* could no doubt have been predicted by sequence analysis, we include it here as to the best of our knowledge, it has not been shown to have a motility defect in *B. subtilis per se*. We note however that the *fliT* mutant flagellar filament defect is entirely consistent with the biochemically-supported role of FliT in *B. subtilis* as a secretion chaperone for the extracellular filament cap FliD (54). The FliT homolog in *Salmonella enterica* is multifunctional serving both as a FliD secretion chaperone and as an adaptor for the regulatory proteolysis of the master activator of flagellar transcription (53,55). Whether *B. subtilis* FliT is also multifunctional remains to be determined but we note that the proteolytic adaptor function (targeting the master activator SwrA) is performed by SmiA, the protein encoded immediately downstream of FliT (7,23,57). Whether *B. subtilis* FliT plays multiple roles in *B. subtilis* flagellar regulation, including the proteolytic modulation of the master activator of flagellar transcription SwrA, is unknown.

Besides the mass confirmation of genes involved in swarming and flagellar biosynthesis, the large TnSeq dataset indicates that the original manual screen used to identify important genes was rather effective as few additional mutants with severe defects were discovered (8). In addition, the system-wide analysis and large dataset allows interpretation of genes that weren’t identified as candidates. For example, the putative “missing parts” in flagellar biosynthesis, specifically the rod polymerization cap protein and peptidoglycan bushings, remain undiscovered. Either *B. subtilis* encodes multiple proteins that are both sequence-divergent and functionally-redundant, or the rod-cap and bushing proteins are unnecessary and should perhaps be considered modifications associated with a Gram negative envelope rather than essential structural components. Likewise, TnSeq revealed no candidate for the peptidoglycan hydrolase activity thought to be necessary for flagellar assembly. While we acknowledge that it remains a possibility that these structural proteins were missed for technical or biological reasons inherent to the screen, we must also acknowledge that there is no evidence *a priori* that such proteins either exist or are necessary in *B. subtilis*.

## MATERIALS AND METHODS

### Strains and growth conditions

*B. subtilis* strains were grown in lysogeny broth (LB) (10 g tryptone, 5 g yeast extract, 5 g NaCl per L) or on LB plates fortified with 1.5% Bacto agar at 37°C. When appropriate, antibiotics were included at the following concentrations: 10 µg/ml tetracycline, 100 µg/ml spectinomycin, 5 µg/ml chloramphenicol, 5 µg/ml kanamycin, and 1 µg/ml erythromycin plus 25 µg/ml lincomycin (*mls*). Isopropyl *β*-D-thiogalactopyranoside (IPTG, Sigma) was added to the medium at the indicated concentration when appropriate.

### Swarm expansion assay

Cells were grown to mid-log phase at 37°C in LB broth and resuspended to 10 OD_600_ in distilled water containing 0.5% India ink (Higgins). Freshly prepared LB containing 0.7% Bacto agar (25 ml/plate) was dried for 10 minutes in a laminar flow hood, centrally inoculated with 10 µl of the cell suspension, dried for another 10 minutes, and incubated at 37°C. The India ink demarks the origin of the colony and the swarm radius was measured relative to the origin. For consistency, an axis was drawn on the back of the plate and swarm radii measurements were taken along this transect. For experiments including IPTG, cells were propagated in broth in the presence of IPTG, and IPTG was included in the swarm agar plates.

### Plate reader growth curves

Colonies were inoculated into 3ml LB broth containing, grown at 37°C in a roller drum until cultures were turbid, and back diluted to an OD_600_ of 0.001. Next, 1ml of each of the cultures were loaded in technical triplicate into a Greiner Bio-One Cellstar 24-well cell culture plate. All remaining wells in the culture plate were filled with 1ml of LB broth, both to serve as a sterile control and to minimize water loss due to dehydration. The plate was loaded onto a BioTek Synergy H1 microplate reader and growth curves were measured using BioTek microplate reader and imager Gen5 3.11 software via the following protocol. Procedure was set as a “kinetic run” for 15 hours with absorbance readings of 600A every 20 minutes. The plate was set to shake in a continuous double-orbital at 37 degrees Celsius with a 1-degree gradient. The procedure was set to read from a Greiner Bio-One Cellstar 24-well cell culture plate with no lid. At the end of the kinetic run the temperature was set to turn off and the plate was held at room-temperature. Data was exported to an Excel spreadsheet and growth rate was calculated over a series of 1 hour windows and the maximum growth rate was recorded.

### Microscopy

Fluorescence microscopy was performed with a Nikon 80i microscope with a phase contrast objective Nikon Plan Apo 100X and an Excite 120 metal halide lamp. FM4-64 was visualized with a C-FL HYQ Texas Red Filter Cube (excitation filter 532-587 nm, barrier filter >590 nm). Alexa Fluor 488 C5 maleimide was visualized using a C-FL HYQ FITC Filter Cube (FITC, excitation filter 460-500 nm, barrier filter 515-550 nm). Images were captured with a Photometrics Coolsnap HQ2 camera in black and white, false colored and superimposed using Metamorph image software.

For fluorescent microscopy of flagella, 0.5 ml of broth culture was harvested at ∼1 OD_600_, and washed once in 1.0 ml of pH 8.0 PBS buffer (137 mM NaCl, 2.7 mM KCl, 10 mM Na_2_HPO_4_, and 2 mM KH_2_PO_4_). The suspension was pelleted, resuspended in 50 µl of PBS containing 5µg/ml Alexa Fluor 488 C_5_ maleimide (Molecular Probes), and incubated for 5 min at room temperature (Blair et al., 2008). 1 ml of PBS was added, cells were pelleted and resuspended in 50 ul of 5 µg/ml FM4-64 (Molecular Probes). Three microliters of suspension were placed on a microscope slide and immobilized with a poly-L-lysine-treated coverslip.

### Strain construction

Mutants were assembled from three primary approaches. First, we used transposon insertions in the gene of interest that were pre-existant in our strain database from past screens. Second, we obtained insertion-deletion alleles in which the kanamycin resistance gene replaced the gene of interest from the BGSC (Bacillus Genetic Stock Center, Columbus OH) originally generated as part of a high-throughput mutagenesis approach (67). Third, we generated our own insertion-deletion alleles with a kanamycin resistance cassette replacement by long-flanking homology with adjacent DNA sequence and Gibson isothermal assembly (57). All strains used in the manuscript are listed in Table S2 and all primers are listed in Table S3

For long flanking homology replacements by isothermal assembly, the regions upstream and downstream of each indicated gene was PCR amplified using the two pairs of primers indicated in the parentheses. Next, DNA containing a kanamycin resistance gene was amplified from plasmid pDG780 (58) was amplified using universal primers 3250/3251. The three DNA fragments were combined at equimolar amounts to a total volume of 5 µL and added to a 15 µl aliquot of prepared master mix (see below). The reaction was incubated for 60 minutes at 50° C. The completed reaction was then PCR amplified using the outside primers to amplify the assembled product. The amplified product was transformed into competent cells of DK1042. Insertions were verified by PCR amplification of mutant chromosomal DNA using the outside primers used for construction. The following primer pairs were used to build the following mutants: *cshA* (6861/6862, 6863/6864); *cshB* (6865/6866, 6867/6866); *dltB* (7175/7176, 7177/7178); *gtaB* (5421/5420, 5419/5418); *rnhC* (7238/7239, 7240/7241); *rnmV* (7179/7180, 7181/7182); *whiA* (7146/7147, 7148/7149); *ykaA* (6876/6877, 6878/ 6879); *ypmA* (7167/7168, 7169/7170); *ypmB* (7142/7143, 7144/7145); *ytlQ* (5769/5770, 5771/5772); and *yvpB* (6853/6854, 6855/6856).

To prepare the isothermal assembly reactions, a 5X isothermal assembly reaction buffer (500 mM Tris-HCL pH 7.5, 50 mM MgCl_2_, 50 mM DTT (Bio-Rad), 31.25 mM PEG-8000 (Fisher Scientific), 5.02 mM NAD (Sigma Aldrich), and 1 mM of each dNTP (New England BioLabs)) was aliquoted and stored at -80° C. An assembly master mixture was made by combining prepared 5X isothermal assembly reaction buffer (131 mM Tris-HCl, 13.1 mM MgCl_2_, 13.1 mM DTT, 8.21 mM PEG-8000, 1.32 mM NAD, and 0.26 mM each dNTP) with Phusion DNA polymerase (New England BioLabs) (0.033 units/µL), T5 exonuclease diluted 1:5 with 5X reaction buffer (New England BioLabs) (0.01 units/µL), Taq DNA ligase (New England BioLabs) (5328 units/µL), and additional dNTPs (267 µM). The master mix was aliquoted as 15 µl and stored at -80°C.

### Transposon library generation and sequencing protocol (TnSeq)

To generate the strain for mutagenesis, DK1042 was transformed with 1 μl of the *mariner* transposon delivery plasmid pWX642 (31) and plated on LB agar containing mls and incubated at 30°C for 48 hours. Four individual colonies were inoculated into one tube containing 4 ml LB spec^100^ broth and rolled at 22°C for 20 hrs and this growth procedure was repeated with 4 parallel tubes. The 4 cultures were pooled, OD_600_ was measured, diluted to OD_600_ = 1 using LB and 500 μl of culture was mixed with glycerol to a final concentration of 14%. Cultures were stored in cryotubes and frozen using liquid nitrogen. To determine the number of transposants per ml, the thawed library was serially diluted, separately plated on LB and LB spec^100^ plates, and incubated overnight at 42°C. The transposition efficiency, determined by dividing the number of spectinomycin resistant colonies by the total number of colonies was approximately 10%.

To generate the libraries, the mutagenized pools were diluted appropriately, plated on 20 large (150 × 15 mm) LB spec^100^ agar plates, and incubated at 42°C overnight such that each plate had ∼50,000 colonies. Colonies were harvested with a rubber scraper, the slurry was diluted to an OD_600_ of 5 and genomic DNA was purified using the DNeasy Blood & Tissue DNA purification kit (QIAGEN). In parallel, 10 μl of glycerol stock was spotted in the center of 10 0.7% LB agar plates and were incubated at 37°C until the swarm edge reached about 25 mm from the origin. The 10 mm edge of the swarm was harvested with a cotton swab into LB media and chromosomal DNA was purified as above from the resulting slurry. The procedure was repeated on five different occasions to produce five motility-enriched libraries.

Next, 200 ng of genomic DNA from each library was digested with MmeI, and each library was separately ligated in barcode tagged adapters with T4 DNA ligase incubated overnight at 16°C. Reactions were PCR amplified, the products were purified and ∼20 ng of each was sequenced on an Illumina NextSeq 500 machine in the Center for Genomics and Bioinformatics (CGB) at Indiana University, Bloomington, IN with a setup of 82 bp single end reads. The reads were trimmed, mapped the *B. subtilis* 3610 genome (NCBI accession CP020102) (59), then analyzed as previously described (25) Genes in which reads were statistically underrepresented in the swarming library compared to the control library were identified by the Mann Whitney U test. Visual inspection of the transposon insertion profiles was performed with the Sanger Artemis Genome Browser and Annotation tool (32).

## ACKNOWLEDGEMENTS

We thank Ankur Dalia and Doug Rusch for the initial TnSeq experiment and analysis, Xheni Karaboja for assisting with TnSeq sample preparation, Zhongqing Ren for helping with TnSeq analysis, the Center for Genomics and Bioinformatics at Indiana University for Illumina sequencing. This work was funded by NIH Grants R01 GM141242 (XW) and R35 GM131783 (DBK).

